# Optimization of Shoot Regeneration and Application of CRISPR/Cas9 Gene Editing to Cultivated Strawberry (*Fragaria* ×*ananassa*)

**DOI:** 10.1101/2023.08.13.553153

**Authors:** Jin-Hee Kim, Cheol-Min Yoo, Chi D Nguyen, Heqiang Huo, Seonghee Lee

## Abstract

Efficient methods of plant transformation and tissue culture are essential to CRISPR/Cas-based gene editing of crops, but neither is well established in cultivated octoploid strawberry (*F* ×*ananassa*). In the present study, a method for shoot regeneration was established and optimized for two strawberry cultivars commercially grown in Florida, Sweet Sensation® ‘Florida 127’ FL127) and ‘Florida Brilliance’ (FB). Runner segments at the tip, node, and petiole obtained from greenhouse-grown plants were used as explants for comparisons of shoot regeneration rate. FL127 showed the highest frequency of shoot regeneration to a basal Murashige and Skoog media (MS) containing 1 mg·L-1 of TDZ, 0.05 mg·L-1 of BA, and 0.05 mg·L-1 of 2,4-D, while FB showed the best response to a lower concentration of BA (0.01 mg·L-1) in the same media type. The average conversion frequencies of somatic embryos into shoot regenerations from the runner tips (RT) were 42.8% in FL127 and 56.9% in FB, respectively, with RT being the most prolific in shoot generation for both cultivars. Using these optimized tissue conditions, *Agrobacterium*-mediated CRISPR/Cas9 gene editing was conducted to evaluate the efficiency of transformation and knockout mutations in the *phytoene desaturase* (*FaPDS*) gene of FL127. A total of 234 explants treated with *Agrobacterium* resulted in an 80.3% regeneration efficiency, with 13.3% of regenerated plants exhibiting partial or complete albino phenotypes. Amplicon sequencing of edited progeny revealed substitutions, insertions, and deletions at the gRNA target sites or flanking regions of all *FaPDS* homoeologous copies. Our results provide effective methods of tissue culture and transformation for the efficient application of CRISPR-mediated gene editing in cultivated strawberry.

## Introduction

The commercial strawberry (*Fragaria* ×*ananassa*) is an allo-octoploid and cultivated worldwide for its popularity and economic value. In the United States, most commercial strawberry cultivars have been developed through conventional breeding methods, recently aided by advanced genomic tools (Whitaker et al. 2020). The technologies of advanced next-generation sequencing (NGS) has paved the way for generating chromosome-scale genome assemblies of polyploid crops and establishing high-throughput genotyping platforms, accelerating the DNA-informed breeding of octoploid strawberry (Edger et al. 2019; Han et al. 2022; Hardigan et al. 2021; Bassil et al. 2015; Verma et al.). Studies have shown that conventional breeding methods combined with marker-assisted and genomic selection can increase selection efficiency and shorten the breeding cycles (Noh et al. 2017; Whitaker 2011; Gezan et al. 2017; Pincot et al. 2020). Additionally, the identification of genes associated with fruit quality and disease resistance in octoploid strawberry has enabled us to utilize genetic engineering techniques for enhancing the breeding traits of new strawberry varieties. With advancements in mutagenesis techniques and plant biotechnology, mutation or transgenic breeding has shifted from random mutagenesis to precise gene editing, allowing for targeted modification of specific gene sequences (Chen et al. 2019). In conjunction with conventional breeding techniques, CRISPR gene editing has the potential to accelerate crop improvement and facilitate the creation of elite plant cultivars through precision breeding.

The recent rapid development of CRISPR/Cas-based gene-editing technology offers new possibilities for crop improvement (Zhang et al. 2020; Biswas et al. 2021; Wolter et al. 2019). Multiple gene-editing approaches have been reported for *Rosacea* crops such as apple, grapevine, pear, and raspberry (Miller 2019; Nishitani et al. 2016; Nakajima et al. 2017; Charrier et al. 2019). In strawberry, the gene-editing has been used to modify auxin biosynthesis genes (*typtophan aminotransferase of Arabidopsis 1* and *auxin response factor 8*) in *F. vesca* and fruit and plant development-related genes (*Tomato MADS box gene6*, *FaTM6*, *phytoene desaturase, PDS,* and *Reduced Anthocyanins in Petioles, RAP*) in *F. ×ananassa* (Zhou et al. 2018; Martín-Pizarro et al. 2019; Wilson et al. 2019; Gao et al. 2020). The mutation of *RAP* gene by CRISPR/Cas9 showed alternation of fruit color both in wild and cultivated strawberry (Gao et al. 2020). The study by Zhou et al (2018) showed that the U6 promoters from either *Arabidopsis* or *F*. *vesca* could effectively drive sgRNA expression in *F. vesca* for CRISPR/Cas9-based gene editing. However, this vector system has not yet been tested in octoploid strawberry. Additionally, CRISPR-based gene editing was performed to mutate *FaPDS* and *FaTM6* in octoploid strawberry, but these studies did not report if the mutations occurred at all possible homoeologous regions of the target gene (Zhou et al. 2018; Martín-Pizarro et al. 2019; Wilson et al. 2019).

The application of CRISPR-mediated gene editing to an octoploid strawberry requires dedicate procedures of tissue culture and the shoot regeneration. There have been successful reports of micropropagation from octoploid strawberry (Rokosa and Mikiciuk 2017; Folta et al. 2006; Nehra et al. 1989; Passey et al. 2003). However, regeneration and morphological development in octoploid strawberry are greatly influenced by tissue culture conditions, explant types, and genotypes (Landi and Mezzetti 2006; Rokosa and Mikiciuk 2017; Debnath and Teixeira da Silva 2007; Folta and Dhingra 2006; Husaini et al. 2011). Therefore, it is often necessary to optimize tissue culture conditions for different strawberry varieties. In this study, an optimal shoot regeneration protocol via indirect somatic embryogenesis from callus generated from runner segments and petioles was established using commercially grown strawberry cultivars, Sweet Sensation® ‘Florida 127’ (FL127) and ‘Florida Brilliance’ (FB). With the optimized conditions, we tested CRISPR-guided mutation of *Phytoene Desaturase* (PDS) gene and determined the mutation efficiency at all homoeologous copies of PDS using the reference genome of FB.

## Materials and Methods

### Plant materials

The runner and petiole segments were collected from four- to six-month-old greenhouse-grown Sweet Sensation^®^ ‘Florida127’ (FL127) and ‘Florida Brilliance’ (FB) plants. For the runners, ∼4 cm long segments adjacent to the basipetal side of the tip (runner tip or RT) and the same size of segments adjacent to the basipetal side of the node (runner internode or RI) were excised and sterilized. For the petioles (P), 4 cm long segments at the leaf end were excised and sterilized (Figure la). Those segments were surface-sterilized by placing them in a diluted solution of household dish soap (0.25%) with continuous stirring for 20 minutes, followed by a bleach solution of 7.5% of Clorox® (0.62% sodium hypochlorite) and 0.25% of Tween 20 with stirring for 20 min. The tissues were rinsed three times with sterile distilled water by soaking five minutes each time, and dried on sterile filter paper (Whatman plc, Maidstone, United Kingdom) in a laminar flow hood. The sterilized tissue segments were trimmed of their bleached ends, then cut into smaller segments (≤ 1 cm) and placed onto tissue culture grade (TC) agar media (PhytoTechnology Laboratories, Lenexa, KS, USA).

### Tissue culture media and conditions

All media used in this study contained mineral salts and vitamins according to Murashige and Skoog (MS) (1962), 3% (w/v) sucrose, plant growth regulators (PGRs), and 0.5% (w/v) of TC agar. The pH was adjusted to 5.75 to 5.8 with KOH before the addition of TC agar. All media were sterilized by autoclaving at 121°C for 15 minutes. To screen for the optimal shoot regeneration media, 56 different media formulations were tested. Each formulation contained 1 mg· L^-1^ of TDZ and a range of 2,4-D concentrations (0, 0.01, 0.02, 0.05, 0.1, 0.2, 0.5 mg·L^-1^), along with varying concentrations of BA (0, 0.01, 0.02, 0.05, 0.1, 0.2, 0.5, 1 mg·L^-1^) (Table 1). The tissues were incubated under cool white fluorescent light at 35 μmol·m^-2^·s^-1^ for 16 hours per day at 25 ± 1°C. After five weeks of growth on the screening media. The media types were evaluated based on the vigor of shoot regeneration, and the best media types were chosen for FL127 and FB and were called SRM (Shoot Regeneration Media) 25 and SRM11, respectively.

**Table 1.** Shoot regeneration media (SRM) types for screening an optimal condition.

| *BA* \ *2,4-D* | *0.00* | *0.01* | *0.02* | *0.05* | *0.1* | *0.2* | *0.5 mg/L* |
| --- | --- | --- | --- | --- | --- | --- | --- |
| *0.00* | 1 | 2 | 3 | 4 | 5 | 6 | 7 |
| *0.01* | 8 | 9 | 10 | 11 | 12 | 13 | 14 |
| *0.02* | 15 | 16 | 17 | 18 | 19 | 20 | 21 |
| *0.05* | 22 | 23 | 24 | 25 | 26 | 27 | 28 |
| *0.1* | 29 | 30 | 31 | 32 | 33 | 34 | 35 |
| *0.2* | 36 | 37 | 38 | 39 | 40 | 41 | 42 |
| *0.5* | 43 | 44 | 45 | 46 | 47 | 48 | 49 |
| *1.0 mg/L* | 50 | 51 | 52 | 53 | 54 | 55 | 56 |

### Regeneration

To optimize shoot regeneration, approximately 100 explants were taken from each of the tissues (RT, RI, and P) of the FL127 and FB plants and cultured on their respective SRMs for four weeks. Subsequently, the whole tissues were sub-cultured to an Elongation Media (EM) to promote shoot differentiation and growth. EM25 for FL127 consisted of 0.05 mg·L^-1^ of IBA, 1 mg mg·L^-1^ of BA, and 100 mg·L^-1^ of proline, while EM11 for FB was made up of 0.01 mg·L^-1^ of IBA, 1 mg mg·L^-1^ of BA and 100 mg· L^-1^ of proline. In these EMs, the shoot clumps grew and expanded over a period of four to six weeks. Once the clumps became loose, they were carefully separated from the callus to promote the growth of smaller clumps. After the shoots elongated to 1-cm in length, the clumps were transferred to hormone-free media (HF) to facilitate rooting. The shoot clumps continued to grow in the HF for forming clusters of plantlets. The rooted individual plantlets were transferred to soil (Metro-Mix®820; Sungro-Horticulture, Agawam, MA, USA) and grown in a growth room under 16-hour light at 25 C°. The number of embryos and shoots generated by each explant was counted in three different types of explants, and the induction levels of callus and shoots were calculated based on the collected numbers. Experiments were repeated three times, and results of each experiment were analyzed using a general linear model and Duncan’s pairwise comparison was conducted to identify significant differences between treatments at the *P* ≥ 0.001 level using SAS software, version 9.2 (SAS Institute, Cary, NC, USA).

### sgRNA design for *FaPDS* and CRISPR/Cas9 vector construction

To validate the efficiency of CRISPR/Cas9 gene editing system under optimized tissue culture conditions, the phytoene desaturase (*PDS*) gene involved in the biosynthesis of chlorophyll, carotenoids, and gibberellins in plants, was selected for study (Qin et al. 2007). The homolog sequence of strawberry *PDS* (*FvPDS*) gene from *F. vesca* (v2.0.a1) was identified from the reference genome of *F. **×**ananassa* in the Genome Database for *Rosaceae* (GDR). Two single-guided (sg) RNAs were designed to target the *FaPDS* gene using CRIPS-P 2.0 (Liu et al. 2017). The previously reported PHN-SpCas9-4xBsaI was used to generate the CRISPR expression construct for targeting all homoeologous copies of *FaPDS* gene (Nguyen et al. 2021). Oligos FaPDSF1 (5’TGCAACAAGCCAGGAGAGTTCAGCGTT-3’) and FaPDSR1 (5’-CTAAAACGCTGAACTCTCCTGGCTTGT-3’) for the *FaPDS* sgRNA1 and *FaPDSF2* (5’-GGTGCACAGAACTTGTTTGGGGAGCT-3’) and *FaPDSR2* (5’-GGTGCACAGAACTTGTTTGGGGAGCT-3’) for the *FaPDS* sgRNA2 were mixed in equal molar mass, denatured at 98°C, and gradually cooled down to room temperature to facilitate oligo annealing. The PHN-SpCas9-4×BsaI was then digested with BsaI and purified using the Wizard® SV Gel and PCR Clean-Up System (Promega, Madison, WI, USA). The annealed oligos FaPDSF1/R1 and FaPDSF2/R2 were mixed and ligated to the purified pHN-SpCas9-4×BsaI to form the final pHN-SpCAS9-FaPDSsgRNAs construct.

### Plant transformation

The *Agrobacterium* co-cultivation method was used for transformation in FL127. The CRISPR/Cas9 vector was transferred to the *Agrobacterium tumefaciens* strain EHA105 through the freeze-thaw transformation method (Weigel and Glazebrook 2006). The EHA105 harboring the binary vector (pHN-SpCAS9-FaPDSsgRNAs) was cultured overnight in a yeast extract peptone (YEP) medium supplemented with Rifampicin (15 mg/L) and Spectinomycin (50 mg/L), and then harvested at 4,200 rpm for 15 minutes using Thermo Sorvall Legend XTR Refrigerated Centrifuge (ThermoFisher Scientific, Waltham, MA, USA). The bacterial pellet was washed and resuspended in a sterilized 1× MS medium containing 0.1mM Acetosyringone and 10mM Glucose (pH 5.8; OD_600_= 0.5). The resuspended *Agrobacterium* was incubated for three hours at room temperature.

Tissue culture explants were prepared following a previously described process. Surface-sterilized runner segments of FL127 were cut into 7 mm-sized segments and mixed with a suspension of the EHA105 strain carrying binary vectors containing Cas9 and two sgRNAs for the target gene. After a 10-min incubation, the co-cultivated segments were transferred to fresh SRM25 medium and incubated in dark conditions for two days. The co-cultivated plant tissues were washed thoroughly with 1× MS medium and cultured on SRM25 containing antibiotics (w/50 mg/L Kanamycin, 100 mg/L Cefotaxime, and 50 mg/L Timentin). The subculture was placed into fresh media every two weeks until plant regeneration, and the albino phenotypes were evaluated.

All explants were subjected under kanamycin selection, and plant regeneration was followed. The success of transformation was validated by checking for the expression of green fluorescent protein (GFP) in embryogenic calli developed in the media containing 50 mg/L of Kanamycin using a fluorescent microscope (Olympus SZX2-ILLT, Olympus Corporation, Tokyo, Japan). Phenotypic analysis of the regenerated transformants was conducted two months after transformation, and the transformants were categorized into five different phenotypes: albino, pale green, light pink, variegated, and green.

### Analysis of CRISPR/Cas9-induced mutation

To determine mutations at the CRISPR/Cas9 target sites of the *PDS* gene, three tissue samples were selected for sequencing analysis. These included two regenerated shoots displaying an albino phenotype and one non-transformed leaf as a wild-type control (FL127). The genomic DNA was extracted from the collected tissues samples following a protocol modified protocol described by Keb-Llanes et al. (2002). The PCR amplicons ranging from 200 bp to 300 bp containing the gRNA target sites were sequenced via Amplicon Sequencing (Genewiz, South Plainfield, New Jersey, USA). Illumina short read were trimmed, aligned, and assembled using Geneious version 11.1.4 (Biomatters Ltd, Auckland, New Zealand) and CLC Genomics Workbench version 11.0.1 (QIAGEN, Aarhus, Denmark). To detect any mutations in the type of SNPs and Indels, contigs with more than 100 short reads from *de novo* assembly were aligned to the high-quality octoploid strawberry reference genome ‘Florida Brilliance’ (FaFB2) (Han et al. 2022; Hardigan et al. 2021).

## Results

### Optimization of media conditions for callus induction and shoot regeneration

The optimal shoot regeneration media (SRM) was screened by culturing explants of three different tissues – runner tip (RT), runner internode (RI), and petiole (P) - on 56 different media types. At the initial stage of tissue culture, the explants induced bright green and compact calli at their cut ends and showed varying degrees of callus growth, with a ranging from no growth to robust growth (3-5mm diameter of calli), depending on the media type. The callus growth was most active in runner tissues as compared to petioles for both FL127 and FB. The callus growth was most active at two weeks of culture as shown in Figure 1b, and was more vigorous at the basipetal end of the runner and petiole tissues. The first shoots were visible after three weeks of culture (Fig. 1b), and after five weeks, the runner and petiole explants showed prolific somatic embryogenesis and shoot induction on two media types for each cultivar (Fig. 1c and 1d). For FL127 the most effective media was a combination of 0.05 mg·L^-1^ of 2,4-D and 0.05 mg·L^-1^ of BA, while for FB it was the combination of 0.05 mg·L^-1^ of 2,4-D and 0.01 mg·L^-1^ of BA. The media types SRM25 and SRM 11 supported a high frequency of shoot regeneration for FL127 and FB, respectively (Table 1).

**Figure 1.**
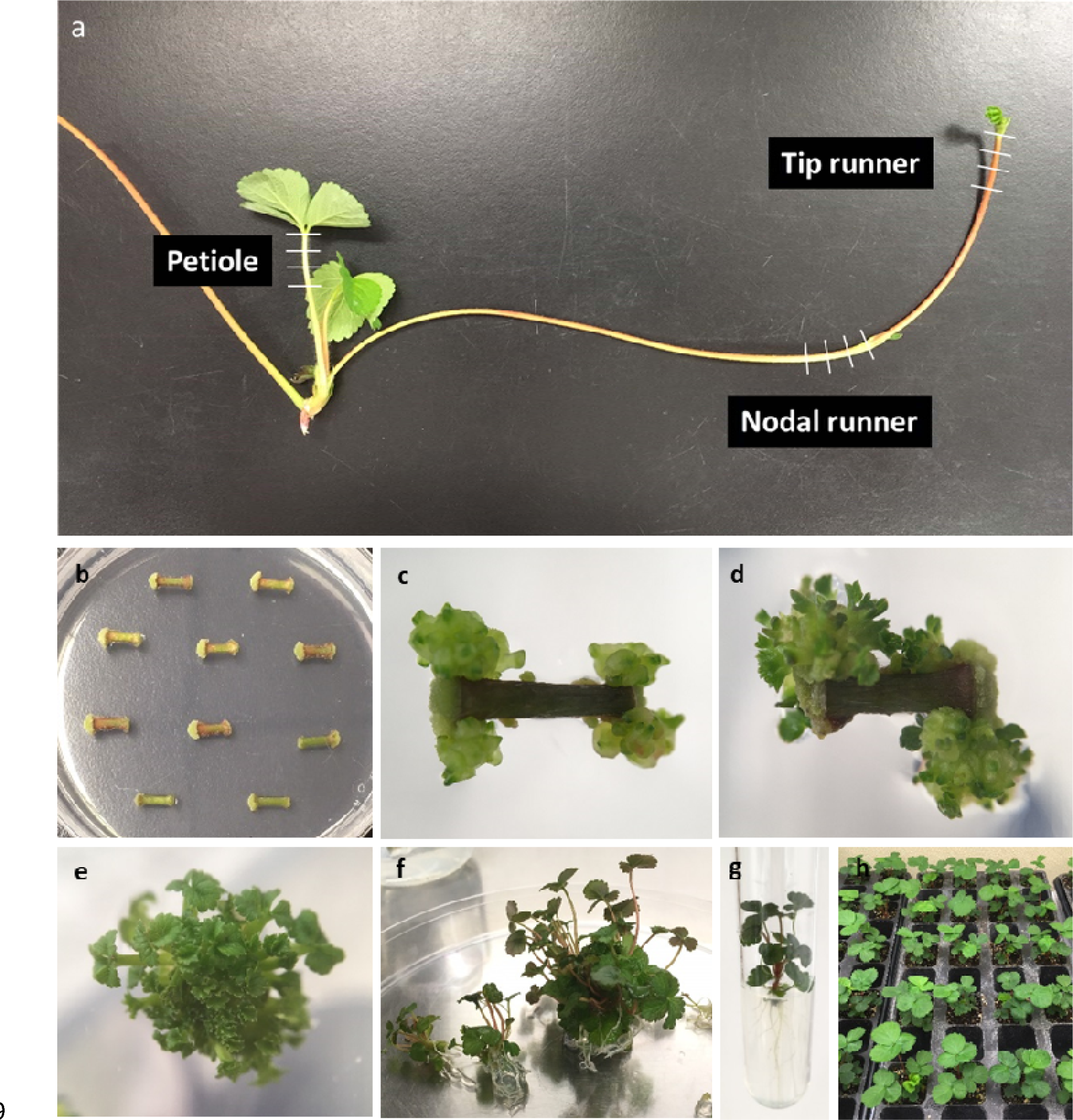
Progression of plant regeneration of Sweet Sensation® ‘Florida 127’. **a** The tissue segments used as the explants. Runner tip (RT), runner internode (RI), and petiole (P) were cut as indicated by the white dashes. **b** Runner segments inducing callus on the Shoot Regeneration Medium. **c** Somatic embryogenesis on the callus on the SRM. **d** Embryos are turning into cotyledonary stage on the Elongation Medium. **e** Isolated shoot clumps growing on the EM. **f** Plantlets are separating from a cluster. **g** An isolated plantlet is growing in a test tube with a hormone-free medium. **h** Plantlets are established in the soil.

### Regeneration of cultivars, FL127 and FB, in optimized tissue culture conditions

All the explants in the optimized SRM successfully induced callus within two weeks as shown in Fig. 1b (Supplementary figure S1a). By the third week of culture, embryos were induced at the edge of the callus with a clear shape indicative of the globular stage of embryos. As early as 18 days into the culture, embryos became visible on the callus from the runner tip tissues in both FL127 and FB, forming tightly held clumps around a solid callus (Fig. 1c and Supplementary figure S1b). In the SRM, the embryos developed into the cotyledonary stage (Fig. 1d and Supplementary figure S1c); however, the embryogenic callus rarely expanded the shoots that matured enough to be converted to plantlets. To promote shoot elongation and maturation, the tissues were sub-cultured in the elongation media, EM25 for FL127 (0.05 mg·L^-1^ of IBA, 1.0 mg· L^-1^ of BA and 100 mg·L^-1^ of proline) and EM11 for FB (0.01 mg·L^-1^ of IBA, 1.0 mg·L^-1^ of BA and 100 mg·L^-1^ of proline). Over a period of four to six weeks of culture in the EMs, the embryos differentiated into shoots, and the subsequent shoot clusters became loose and detached from the callus (Fig. 1e and Supplementary figure S1d). At this stage, the number of shoots per explant was counted. The expanded shoot clusters were transferred to hormone-free media (HF) for rooting, and roots were induced from the shoot clusters, resulting in a bunch of independent plantlets that could be detached from the cluster over two to four weeks. The isolated plantlets grown in HF media were moved to soil (Fig. 1f, g, h, and Supplementary figure S1e). The whole process from placing tissues in media to transferring into soil is about 14 to 16 weeks.

### Tissue specificity for shoot regeneration

Among the three different explants, the runner tip (RT) was the most prolific for callus induction and subsequent shoot regeneration in both FL127 and FB. Callus formation on RT was observed as early as three days of culture, while it took five to seven days for runner internode (RI), and petiole (P). The frequency of embryo induction was highest in RT with 81.4% in FL127 and 84.2% in FB. On the other hand, the lowest percentage of embryogenesis was observed in P (11.7% of explants in FL127 and 59.1% in FB), which RI showed an intermediate percentage (32.1% in FL127 and 14.0% of FB) (Table 2). Although RT showed the highest number of embryos per explant in both cultivars, the difference was not as pronounced as the embryogenesis frequency among the explants. The average number of shoots per explant was counted after the embryo developed into shoots at the later stage in EM (Fig. 1d). The conversion frequency of the embryos was obtained by dividing the number of shoots by the number of embryos. Interestingly, P from FL127 showed the highest conversion rate of embryos to shoots (70.7%), despite inducing the least number of embryos (11.7 ± 1.6 embryos per explant). On the other hand, RT had the lowest conversion rate (42.8%), but it induced the highest number of embryos (81.4 ± 2.0) among the explants for FL127. Overall, RT was the most prolific in shoot regeneration and produced the highest number of regenerated shoots in both cultivars (Table 2).

**Table 2.**
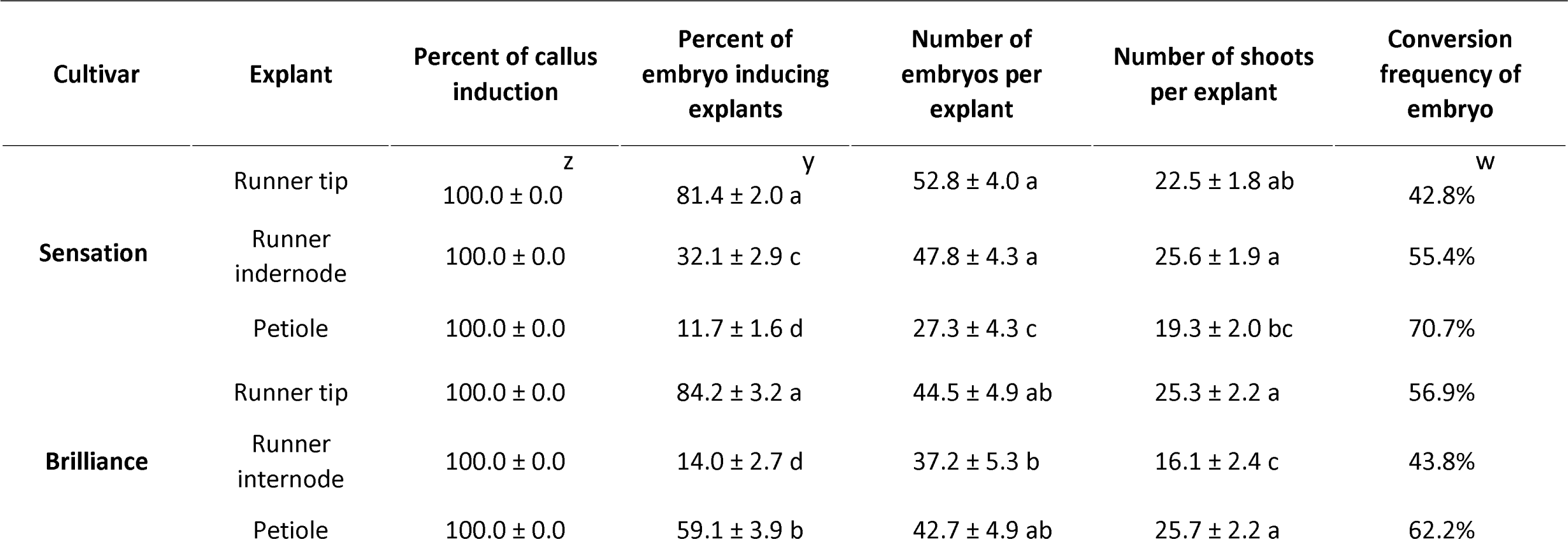
Effect of different explants of the strawberry plants on callus induction, somatic embryogenesis, and shoot regeneration in their optimal media condition.

| Cultivar | Explant | Percent of callus induction<sup>z</sup> | Percent of embryo inducing explants<sup>y</sup> | Number of embryos per explant | Number of shoots per explant | Conversion frequency of embryo<sup>w</sup> |
| --- | --- | --- | --- | --- | --- | --- |
| **Sensation** | Runner tip | 100.0 ± 0.0 | 81.4 ± 2.0 a | 52.8 ± 4.0 a | 22.5 ± 1.8 ab | 42.8% |
|  | Runner internode | 100.0 ± 0.0 | 32.1 ± 2.9 c | 47.8 ± 4.3 a | 25.6 ± 1.9 a | 55.4% |
|  | Petiole | 100.0 ± 0.0 | 11.7 ± 1.6 d | 27.3 ± 4.3 c | 19.3 ± 2.0 bc | 70.7% |
| **Brilliance** | Runner tip | 100.0 ± 0.0 | 84.2 ± 3.2 a | 44.5 ± 4.9 ab | 25.3 ± 2.2 a | 56.9% |
|  | Runner internode | 100.0 ± 0.0 | 14.0 ± 2.7 d | 37.2 ± 5.3 b | 16.1 ± 2.4 c | 43.8% |
|  | Petiole | 100.0 ± 0.0 | 59.1 ± 3.9 b | 42.7 ± 4.9 ab | 25.7 ± 2.2 a | 62.2% |

### Application of CRISPR/Cas9 gene editing in Sweet Sensation^®^ ‘Florida 127’

CRISPR/Cas9 gene editing system was utilized in the optimized condition of tissue culture and shoot regeneration to target a *phytoene desaturase* (*PDS*) gene. This gene was selected due to its role in causing distinct dwarf and albino phenotypes in loss-of-function mutants. A single *PDS* gene was identified on chromosome 4 in the diploid genome of *F*. *vesca*, ‘Hawaiian 4’ (XM_004296916.2) with a sequence similarity of approximately 93% to the *FaPDS* gene from octoploid strawberry (*F.* ***×****ananassa*) (Fig. 2a). The four homoeologous copies of *FaPDS* in *F.* ***×****ananassa* were found to be nearly identical with only a few SNPs, and all sequences from both *F. vesca* and *F.* ***×****ananassa* had the same length at 569 amino acids and 14 exons (Fig. 2a).

**Figure 2.**
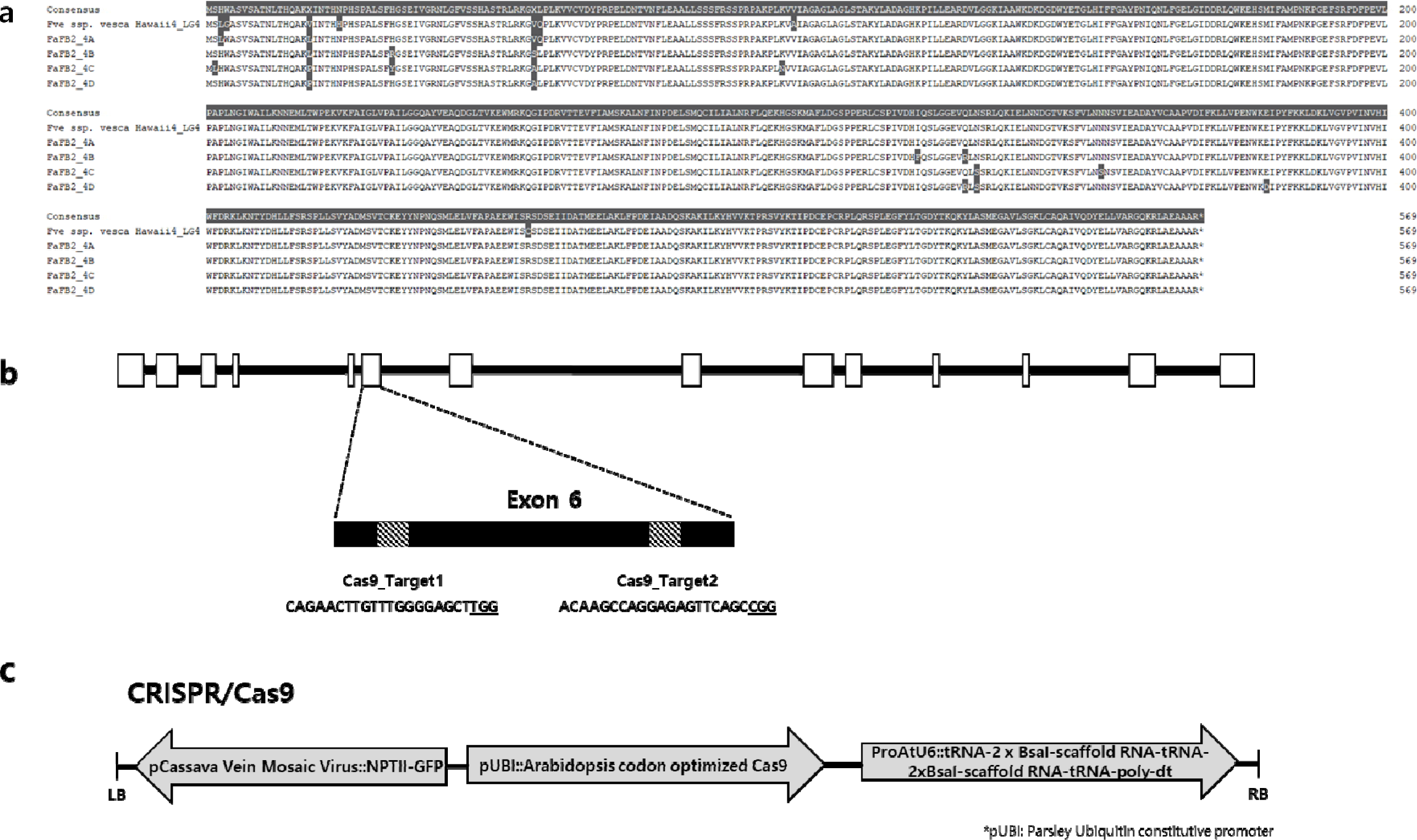
a. Amino acid sequences of *phytoene desaturase* from *Fragaria vesca* and FaFB2. **b** Target sites of CRISPR/Cas9 in *FaPDS* gene. White boxes represent exons of *FaPDS* and the two target sites for CRISPR/Cas9 are located at Exon 6. **c** Binary vector map of CRISPR/Cas9. pHN-SpCas9-4xBsaI, containing a fused NPTII-GFP driven by the Cassava vein mosaic virus promoter for antibiotic and fluorescence selection and a ProAtU6-26-tRNA-2×BsaI-scaffold RNA-tRNA-2×BsaI scaffold RNA polydT cassette

To design the CRISPR/Cas9 construct, two 20-bp sequences containing the 5’-NGG-3’ PAM site were selected. The target sites of CRISPR/Cas9 (Cas9-Target1 and Cas9-Target2) were located at the sixth exon of *FaPDS* (Fig. 2b). Two CRISPR/Cas9 target sequences (5’-CAGAACTTGTTTGGGGAGCT<u>TGG</u>-3’ and 5’-ACAAGCCAGGAGAGTTCAGC<u>CGG</u>-3’) were identical to the sequences of *PDS* genes in both the diploid *F. vesca* and octoploid *F*. ×*ananassa* genomes. To verify the specificity of the off-target sequences, 20-bp target sequences were blasted to the reference genomes of *F. vesca* and *F*. ×*ananassa* (‘Royal Royce’ - FaRR1 and ‘Florida Brilliance’ - FaFB2). It was confirmed that the target sgRNA sequences were specific to all homoeologous copies of *FaPDS* in chromosome 4 and inserted into pHN-SpCAS9-FaPDSsgRNAs vector for CRISPR-mediated gene knockout (Fig. 2c).

### Efficiency of *Agrobacterium*-mediated transformation and CRISPR/Cas9-induced mutation of *FaPDS* gene

A total of 234 strawberry explants from P and RT were used for *Agrobacterium*-mediated transformation and CRISPR/Cas9 gene editing in FL127. The efficiency of regeneration from *Agrobacterium*-mediated transformation for all explants was 80.3%. The 25 tissues (13.3% of the regenerated tissues) showed potential *FaPDS* knock-out phenotypes (Table 3). Corresponding to the previous embryogenesis result, the higher embryogenesis rate was shown in RT (93.3%) compared to P (63.0%) in the FL127 plants. Green fluorescent protein (GFP) signal in the explants was monitored five days after transformation to confirm transformed embryos. GFP spots were observed in most of the transformed tissues (Fig. 3a and b). However, the GFP spots were not clearly visible as the embryos differentiated organs, such as shoots or leaves. Furthermore, only a few GFP-positive embryos developed shoots or other organs. As the embryos differentiated, the transformed shoots and leaves were phenotyped two months after transformation. The phenotypes of the *FaPDS* mutants were shown as albino, pale green, pink, and variegated (Fig. 3c-e). A total of 25 *FaPDS* mutants were confirmed, each with different phenotypes and plant regeneration stages (Table 3). Most of the *FaPDS* mutants showed light pink phenotypes with pink edges and bleached pink centers on the leaves (Fig. 3e). Only a few shoots with albino phenotype were found (Fig. 3c).

**Figure 3.**
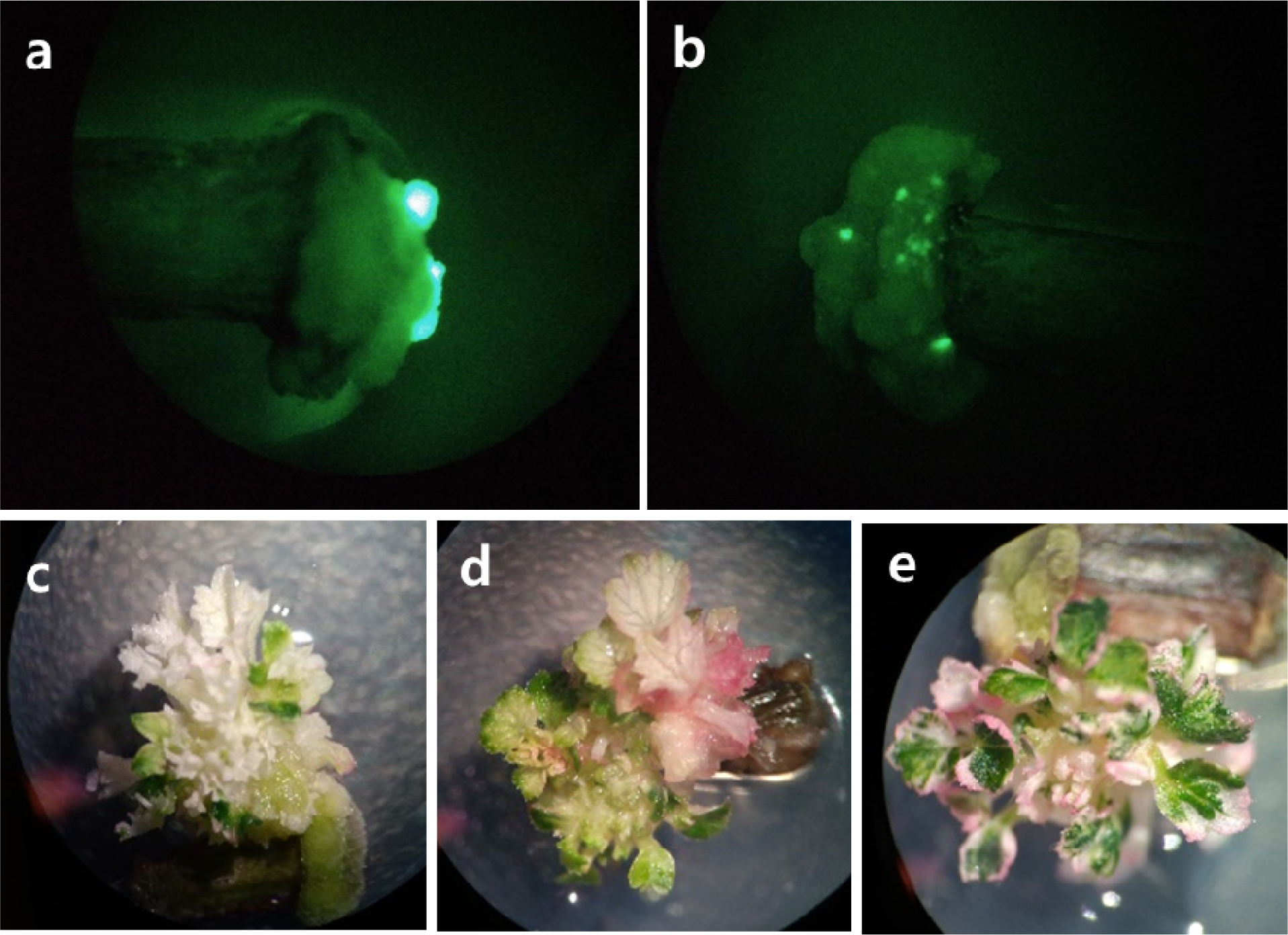
Phenotypes of *FaPDS* transformed shoots and leaves induced by CRISPR/Cas9. **a b** GFP signals were monitored on transformants a week after transformation. Clear albino (**c**), light pink (**d**), and variegated (**e**) transformants were observed two months after transformation.

**Table 3.** The list of collected samples from *FaPDS* transformants induced by CRISPR/Cas9.

| **Regenerants** | **Phenotype** | **Tissue Stage** |
| --- | --- | --- |
| Cas9-1 | white | embryos |
| Cas9-2 | mix (pink, white, green) | embryos |
| Cas9-3 | green | embryos |
| Cas9-4 | mix (white, green) | shoots |
| Cas9-5 | white | embryos |
| Cas9-6 | white | embryos |
| Cas9-7 | white | embryos |
| Cas9-8 | green | shoots |
| Cas9-9 | mix (green, pink) | shoots |
| Cas9-10 | white | shoots |
| Cas9-11 | white | shoots |
| Cas9-12 | green | shoots |
| Cas9-13 | white | callus |
| Cas9-14 | mix (white, green) | embryos |
| Cas9-15 | white | embryos |
| Cas9-16 | pale yellow | shoots |
| Cas9-17 | white w/ pink edge | shoots |
| Cas9-18 | pale green | embryos |
| Cas9-19 | white | embryos, callus |
| Cas9-20 | white | embryos |
| Cas9-21 | mix (pink, white, green) | shoots |
| Cas9-22 | white | embryos |
| Cas9-23 | pinky white | embryos |
| Cas9-24 | green | shoots |
| Cas9-25 | white | shoots |

To examine the presence of mutations at the target sites, two *FaPDS* mutants and one non-transformed were analyzed by Amplicon sequencing (Genewiz, South Plainfield, New Jersey, USA). The 833-bp PCR products containing the target sgRNA site were sequenced and mapped to the haplotype-phased reference genome of FaFB2 (Han et al. 2022; Hardigan et al. 2021). Different types of mutations, including substitutions, insertions, and deletions, were detected on the target sites or flanking regions. The *de novo* assembly result of Cas9-DNA10 revealed two large deletions (CRISPR/Cas9-DNA10_M1 and M2) at the CRISPR/Cas9-Target 1 site. By determining subgenome specific SNPs present at the flanking regions of the deletion, it was determined the Cas9-DNA10_M1 deletion was in chromosome 4A and Cas9-DNA10_M2 in chromosome 4D (Fig. 4a). In addition, a single nucleotide change was found within the CRISPR/Cas9-Target 2 site (CRISPR/Cas9-DNA10_M3, Fig. 4a). The 19th nucleotide guanine (G) was substituted with adenine (A), resulting in a synonymous substitution in chromosome 4C. the assembly result of mutant Cas9-DNA25 showed one SNP change within the CRISPR/Cas9-Target 1 and a 24-bp deletion between Target 1 and 2 (Fig. 4a). The third nucleotide guanine (G) in the Cas9-Target 1 site was substituted with thymine (T), resulting in a synonymous mutation. Additionally, one substitution from cytosine (C) to adenine (A) was found in front of the Cas9-Target 2 site, which could result in a nonsynonymous change, and was found at chromosome 4B. A 24-bp deletion was found at chromosome 4A. Our findings indicate that different types of mutations were observed in all homoeologous genomic regions of *FaPDS* that could contribute phenotypic changes.

**Figure 4.**
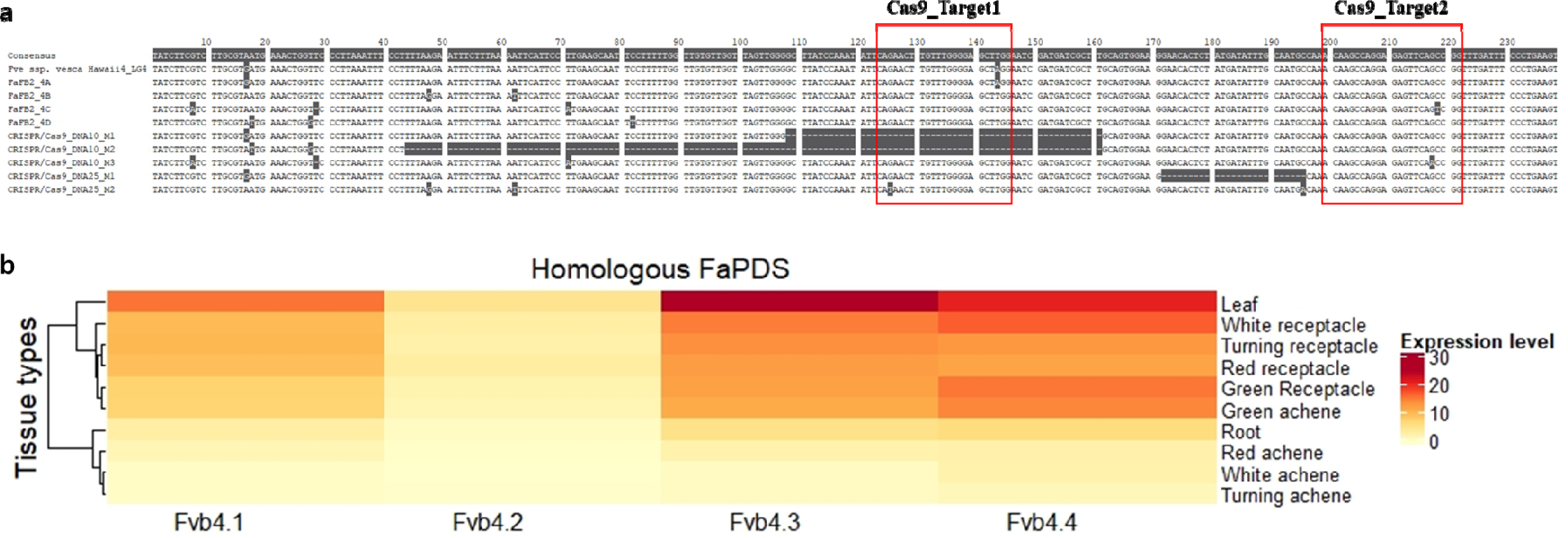
Mutations in the *FaPDS* gene induced by CRISPR/Cas9. **a** Mutant DNA10 and DNA25 were aligned with reference genomes; *F. vesca spp. vesca* Hawaii4, *F.* ***×****ananassa* FaRR1, and *F.* ***×****ananassa* FaFB2. The red boxes indicate the two target sites of CRISPR/Cas9. The nucleotides marked with dark grey indicate single nucleotide polymorphisms. Hyphens within sequences show deleted sequences by CRISPR/Cas9 genome editing. **b** Gene expression pattern of homologous *FaPDS* genes in 10 different tissue types in cultivated strawberry plant (*F.* ***×****ananassa* ‘Camarosa’).

## Discussion

Octoploid strawberry (*F.* ***×****ananassa*) is a highly heterogeneous and diverse species. The success of *in vitro* plant transformation and regeneration greatly depends on the availability of optimized tissue culture conditions. In addition, the transformation efficiency and shoot regeneration rate can vary greatly among cultivars (Folta and Dhingra 2006; Landi and Mezzetti 2006; Passey et al. 2003). Most of shoot regeneration protocols developed in octoploid strawberry are devised using leaf or petiole tissues, while somatic embryogenesis has been induced using different types of tissues such as leaves, petioles, flower buds, and immature embryos (Biswas et al. 2007; Donnoli et al. 2001; Husaini and Abdin 2007; Kordestani and Karami 2008; Pallavi et al. 2011; Lis ; Wang et al. 1984). Indirect somatic embryogenesis from greenhouse-grown petioles was reported in ‘Sweet Charlie’, which is one of the ancestral parents of FL127 (Pallavi et al. 2011; Whitaker et al. 2015). In this study, petiole tissues from FL127 were used for tissue culture with the same concentration of TDZ and 2,4-D used in ‘Sweet Charlie’. This suggests that tissue culture media and conditions may be highly depend on their genetic relationships between cultivars in octoploid strawberry.

The runner segment is not a commonly used explant for strawberry micropropagation (Rokosa and Mikiciuk 2017). This is because most regeneration protocols were developed using *in vitro* cultured plantlets. Only two studies have reported shoot regeneration from runner segments in ‘Allstar’, ‘Honeoye’ and ‘LF9’ strawberry cultivars (Liu and Sanford 1988; Folta et al. 2006). Both studies showed lower frequencies of shoot organogenesis from runner tissues. However, in this study, we successfully achieved an efficient indirect somatic embryogenesis from runner tissues of FL127 and FB. It was often found that the somatic embryos developed not only from the callus at the end of segments, but also at the shank of the runner or petiole tissues (Supplementary Fig. S2a). It was also consistently observed that when there were a large number of embryos on a callus, the embryos often remained at the heart or torpedo-shaped stage without producing shoots. This may be due to competition for resources among the embryos and could have contributed to the lower conversion frequency of embryos on explants with more embryos (Table 2). TDZ was used as the main growth regulator in the SRMs, as it has been found to substitute for both auxin and cytokinin in many species and is highly effective at inducing somatic embryos (Landi and Mezzetti 2006; Cappelletti et al. 2016; Mithila et al. 2003). However, TDZ can inhibit shoot elongation and differentiation (Ouyang et al. 2016; Huetteman and Preece 1993). When TDZ remained in the SRMs, the somatic embryos developed to the cotyledonary stage but did not grow further into shoot clusters. Thus, it was critical to subculture the embryo-generating explants from the SRM to the EM by the fifth week of culture.

The feasibility of CRISPR/Cas gene-editing in octoploid strawberry has been evaluated by recent studies (Zhou et al. 2018; Martín-Pizarro et al. 2019; Wilson et al. 2019; Zhu et al. 2020). In this study, we optimized tissue culture conditions for two major cultivars grown in Florida, Sweet Sensation® ‘FL127’ and ‘Florida Brilliance’ and used these conditions to determine the efficiency of CRISPR/Cas9 vector constructs and mutations through *Agrobacterium*-transformation. Using the recently published chromosome-scale haplotype-phased genome of FaFB2, we were able to identify all homeologous copies of the *FaPDS* gene on chromosome 4 (Han et al. 2022). It has been known that one subgenome (*F. vesca*-like) is dominant to others in octoploid strawberry (Hardigan et al. 2020). As shown in Figure 4b, our analysis of gene expression patterns using transcriptome data from 10 different tissues of ‘Camarosa’, showed that all four homoeologous copies of *FaPDS* are expressed in the receptacle, leaf, and green achene, but have low expression in the root, white, turning, and red achenes (Barbey et al. 2020; Sánchez-Sevilla et al. 2017). The copy located at chromosome 4-3 (4A in FB) showed the highest level of expression in green leaf tissues, while the copy at chromosome 4-2 (4C in FB) showed low expression levels. These results indicate that three homoeologous *FaPDS* could be functionally expressed in different parts of strawberry tissues.

In recent studies of CRISPR/Cas9 gene editing in strawberries, the rates of mutant phenotypes produced through *Agrobacterium*-mediated transformation has been found to vary widely due to factors such as promoter types, ploidy levels, and target sequences (Zhou et al. 2018; Xing et al. 2018; Wilson et al. 2019; Martín-Pizarro et al. 2019). In our study, 188 out of 234 transformed explants (P and RT; 80.3%) produced embryos through *Agrobacterium*-mediated CRISPR/Cas9, and 25 transformants were collected. However, we found that most of regenerated explants were not effectively selected using kanamycin, a commonly used selection agent for plant transformation at a concentration of 50 mg/L (Martín-Pizarro et al. 2019; Nehra et al. 1990; Estopà et al. 2001). According to Wilson et al. (2019), the concentrations of kanamycin used for selection may vary for different tissue types in strawberry. Approximately 46% of putative transformants showed the *FaPDS* mutant phenotype after the regeneration of shoots on kanamycin selection medium (Wilson et al. 2019). It would be essential to test kanamycin concentrations in different cultivars to maximize the efficiency of transformation. In addition, another selection reagent such as hygromycin could be tested for optimal concentrations and used for enhancing the transformation rate of CRISPR gene editing in cultivated strawberry.

In conclusion, our study presents regeneration protocols for octoploid strawberry cultivars Sweet Sensation® ‘Florida127’ and ‘Florida Brilliance’ and demonstrates the application of the CRISPR/Cas9 gene editing system. Our results indicate that CRISPR-mediated mutation occurred in one and/or multiple homoeologous copies of the *FaPDS* gene. The tissue culture and regeneration system developed in this study can be easily modified for use in different strawberry cultivars. These results were shown that runner explants from greenhouse plants can be also favorable to use for tissue culture and gene editing with high regeneration rates instead to explants from in vitro plants. Moreover, our CRISPR gene editing results were evaluated with the recently released haplotype-phased genome, and the mutations were examined at the subchromosomal level. Therefore, we confirmed the mutations were found in all *PDS* copies located in four different subchromosomes. With recent advances in the discovery of QTLs and candidate genes controlling important traits, the optimized tissue culture and CRISPR gene editing system in octoploid strawberry can be efficiently utilized for cultivar improvement and functional genomics studies in strawberry.

## Abbreviations

TDZ: *N*-phenyl-*N’*-1,2,3-thiadiazol-5-ylurea
BA: *N*^6^-enzyladenine
2,4-D: 2,4-dichlorophenoxy acetic acid
IBA: indol-3-butyric acid
CRISPR: clustered regularly interspaced short palindromic repeats
Cas9: CRISPR/associated endonuclease
PDS: *phytoene desaturase* gene
NGS: next-generation sequencing.

## Notes

### Competing Interest Statement

The authors have declared no competing interest.

